# Elevated carbon dioxide stimulates highly efficient organic-carbon consumption and confectionary-waste valorization under mixotrophy in the unicellular alga *Galdieria*

**DOI:** 10.1101/2025.05.22.655468

**Authors:** Mauricio Lopez Portillo Masson, Bárbara Bastos de Freitas, Andrei Zybinskii, Ghalih Althagafi, Ma’an Amad, Michael D. Fox, Peter J. Lammers, Kyle J. Lauersen

## Abstract

Unicellular algae are appealing for nutritional and biotechnological utility but have wide variation across strains and can be challenging to produce. The thermo-acidophilic algal genus *Galdieria* use diverse organic-carbon sources for fermentative growth that can include waste-stream feedstocks and have complete amino-acid compositions for human nutrition. Here, we investigated *Galdieria* metabolic dynamics to catalog organic-carbon conversion to biomass. Tested strains had enhanced growth upon 3% CO_2_ supplementation, triggering efficient glucose uptake to reach ∼5 ± 0.3 g dry biomass L^⁻¹^. Stable-isotope analysis revealed that organic-carbon uptake dominates CO₂ fixation in darkness under mixotrophy, with CO₂ an apparent metabolic trigger. *Galdieria sulphuraria* 5587.1 can consume up to 8.3 g carbon L^−1^ day^−1^ from industrial confectionery waste, with C-phycocyanin reaching 3.8% of dry biomass and remaining thermostable at 72°C. This framework can optimize *Galdieria*-based bioprocesses for inexpensive waste conversion into high-value biomass and identifies CO₂ as a trigger of organic-carbon assimilation, even in heterotrophic conditions.

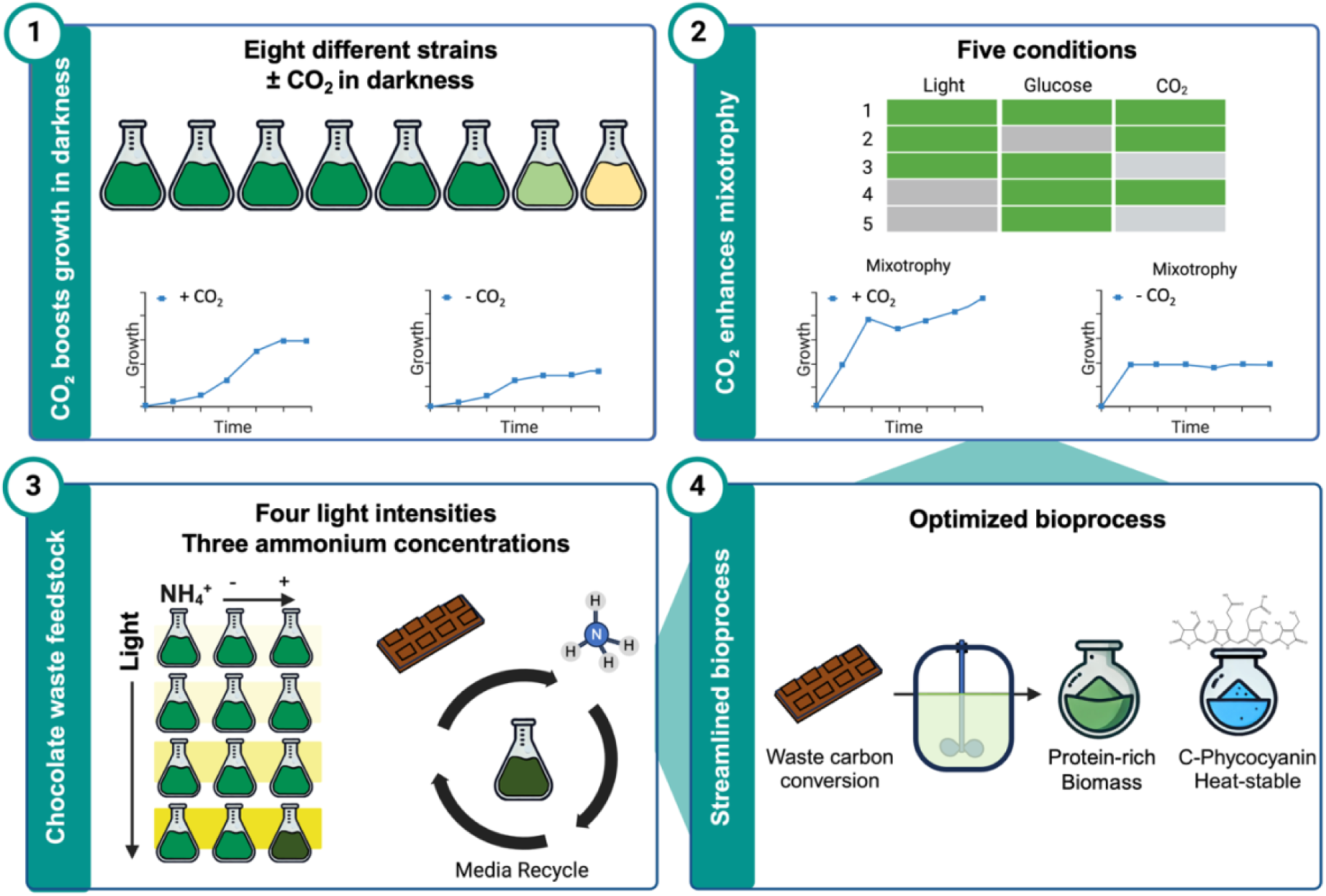

## Introduction

The Cyanidiophyceae are an ancient class of unicellular red algae that inhabit acidic hot springs globally. They exhibit tolerance to temperatures up to 56°C, very low pH, as well as high concentrations of metals [1]. Some have even been found on tree stumps in industrial coal-waste heaps, where biogenic processes and subterranean combustion lead to elevated temperatures [2]. This class currently contains four Genera: *Cavernulicola*, *Cyanidium*, *Cyanidioschyzon*, and *Galdieria* [3], with the latter two being the most well-studied [4]. *Galdieria* was first described in 1981 and exhibits a complex morphology with a tough cell wall [5]. *Galdieria* maintain low GC contents in their nuclear genomes (∼45%), a diploid vegetative state, and can consume 34 known carbon sources [6–10]. Their sexual cycle was recently described, whereby pH controls the conversion of diploid cells to haploids lacking cell walls, and in this state, genetic transformation via homologous recombination as well as classical mating were achieved [11].

All Cyanidiophyceae retain the ancestral cyanobacterial light harvesting complex pigment C-phycocyanin (C-PC) within their plastids [12]. The capacity for organic-carbon consumption led to the proposed use of *Galdieria* species in fermentative biotechnological processes aimed at the conversion of carbon sources into its biomass containing the economically valuable blue water-soluble C-phycocyanin (C-PC) pigment [7,12–15]. Natural C-PC has become increasingly important in the food, pharmaceutical, and nutraceutical industries. It is used as a coloring in various types of candies, drinks, and dairy products. It is regarded as a sustainable alternative to synthetic dyes. C-PC also has antioxidant and anti-inflammatory properties, making it fit for use in dietary supplements and therapeutic products. The market for phycocyanin is projected to grow to USD 276.4 million by 2030 and pharmaceutical-grade C-PC market alone was valued at ca. USD 31.5 million in 2024 (https://www.grandviewresearch.com/industry-analysis/phycocyanin-market-report, https://www.globalgrowthinsights.com/market-reports/pharmaceutical-grade-phycocyanin-market-101222?utm).

In addition to glucose, *Galdieria* are capable of consuming many carbon sources that are considered waste products, like glycerol, xylose, and xylitol [6]. The amino-acid content of *Galdieria* biomass is closer to animal meats than plant-based proteins, a feature that has led to the proposal of novel food status in the European Union (<u>see Fermentalg application</u> 2019 and [16,17]). However, under organic-carbon-feeding regimes, certain strains lose pigmentation and photosynthetic capacity, contrary to the specified goals of producing C-PC. Different *Galdieria* species and strains from around the world vary in pigment contents under organic-carbon consumption, however, dynamics of C-PC accumulation in these strains have not been well described [3,16,18,19].

Strains of *Galdieria* that can maintain pigmentation while growing with organic carbon are capable of mixotrophy: the simultaneous consumption of organic carbon and inorganic carbon fixation through the Calvin–Benson–Bassham (CBB) cycle [20]. Understanding their cultivation dynamics and carbon-accumulation abilities is important for proposing scaled bioprocesses. Various reports have demonstrated yields of *Galdieria* biomass in illuminated mixotrophic-cultivation setups on different organic-carbon substrates: wastewater [21–23], food waste hydrolysates [24], or glucose [17,25–27]. Glucose-fed cultures using dissolved oxygen to balance organic-carbon feeding in oxygen-balanced mixotrophy (OBM) have very high biomass yields [26,27].

Industrial food production wastes also present an abundant yet under-used resource that could be repurposed for algae production, particularly using extremophilic organisms such as *Galdieria*. The chocolate industry generates significant amounts of organic waste, estimated at ∼2% dry mass of ingredients, including chocolate mass and butter, that are discarded with no other use-case during confectionery production [28,29]. These by-products are rich in fermentable sugars, fats, and proteins [28], making them suitable substrates for microorganisms. *Galdieria* have strong potential for metabolizing such complex organic waste streams, efficiently converting them into protein-rich biomass while maintaining pigment production [7,9,13,21,24]. Additionally, its acidophilic and thermotolerant nature enables cultivation in non-sterile conditions, reducing the risk of contamination and lowering processing costs. Because *Galdieria* can thrive in extreme conditions, it has a unique advantage in waste valorization processes since growth of other microorganisms can be suppressed. Food-industry waste is a cost-effective carbon source for *Galdieria* cultivation, repurposing it into high-value bioproducts like C-PC [24]. We were interested in understanding the dynamics of *Galdieria* in different growth modes, and whether strains with the stay-green-in-glucose phenotype could be exploited to effectively convert waste carbon to biomass. We found several surprising features of *Galdieria* behavior in this process, including how organic carbon is favored only under high CO_2_, its amenability for growth in recycled media supplemented with commercial confectionary waste, and how an economically important pigment has enhanced thermostability and production yield than competing commercially used organisms. Our results form an important matrix of understanding to frame future implementation of mixotrophic or heterotrophic processes that seek to convert wasted carbon into protein-rich and pigment-containing *Galdieria* biomasses. This is the first report pinpointing CO_2_ as a metabolic trigger for organic-carbon consumption, even during heterotrophic processes.

## Results & Discussion

### Fv/Fm is reduced in presence of organic carbon across the *Galdieria* genus

As fermentable algae, *Galdieria* species have been investigated for their broad carbon-substrate range [6,13]. We set out to investigate the relationship between organic-carbon use and mixotrophy in this genus, with the aim to find ideal conditions for its biomass production and C-PC accumulation. This was prompted by high carbon conversion to biomass yields reported using oxygen-balanced mixotrophy (OBM) [26,27]. Here, seven strains of *G. sulphuraria* and *G. partita* N1 isolated from different global locations were cultivated for 8 days in heterotrophic conditions to first investigate their pigment-retention behaviors using glucose as a sole carbon source in darkness, with or without 3% CO_2_ supplementation (Figure 1A). Strains *G. partita*, *G. sulphuraria* 5587.1, Soos, and 074G exhibited a dark-green culture phenotype in these conditions (Figure 1A). However, *G. sulphuraria* 5572 and 3377 displayed lighter green phenotypes, while 21.92 and 074W lost pigmentation completely (Figure 1A). Regardless of pigmentation or elevated CO_2_, photosystem II linear electron-transfer rates (Fv/Fm) were reduced during cultivation, with 21.92 and 074W showing almost no detectable activity concomitant with pigment loss (lower panel). Strains that maintained dark pigmentation had stable but reduced Fv/Fm rates up to day 8. Variation in pigmentation loss and stabilization of Fv/Fm suggest that certain strains (*G. partita* N1, *G. sulphuraria* 5587.1, Soos, and 074G) possess metabolic adaptations that enable at least partial retention of photosynthetic machinery. Supplementation of 3% CO_₂_ did not prevent the decline in Fv/Fm in any strain, separating this behavior from the presence of CO_2_.

**Figure 1.**
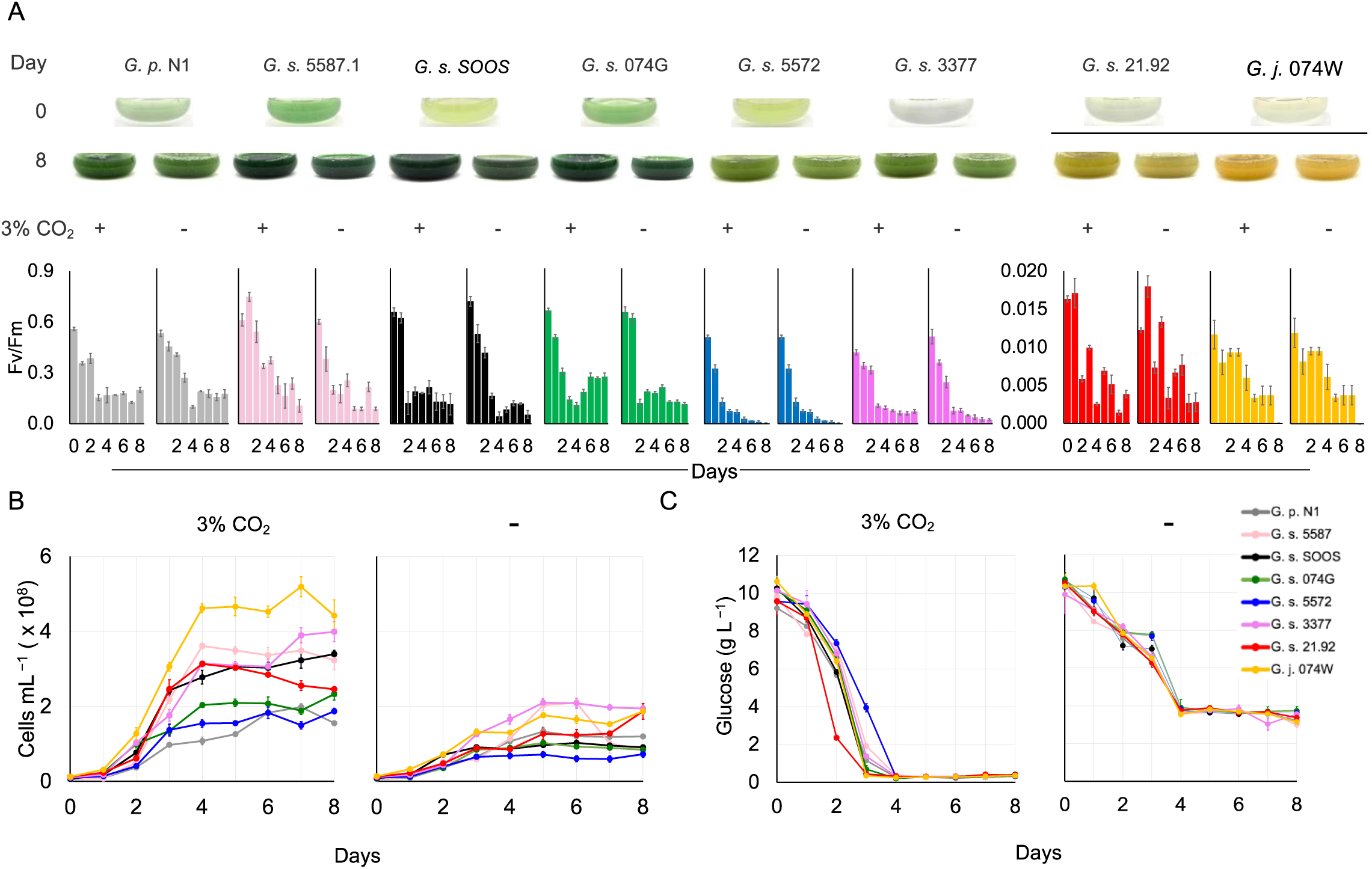
Impact of organic carbon and CO_₂_ supplementation on pigment retention and metabolism in *Galdieria* strains. **A**. Phenotypic variation of eight *Galdieria* strains cultivated in MA2 media with glucose 10 g L^⁻¹^ as the sole carbon source in the dark, with or without 3% CO_₂_ atmosphere. Representative images of cultures at T_0_ and after 8 d of cultivation in 400-mL bioreactors are shown. Lower panel shows Photosystem II efficiency (Fv/Fm) during heterotrophic growth of cultures as in. **B, C.** Growth dynamics (B) and glucose consumption (C) by *Galdieria* strains grown under heterotrophic conditions. Error bars indicate the standard deviation of n = 3 replicates.

### Sugar consumption by *Galdieria* stalls without elevated CO_2_

We next investigated the behaviors of all seven *Galdieria* strains in 8-d heterotrophic batch cultivations using 10 g L^−1^ glucose ± 3% CO_2_ (Figure 1B,C). Cell density was higher, and glucose consumption complete, only in cultures supplied with 3% CO_2_ (Figure 1B,C). Cultures without CO_2_ supplementation stalled glucose consumption at ∼4 g L^−1^ (Figure 1B). Similar behavior was reported in larger mixotrophic cultures of *G. sulphuraria* ACUF 064 [26]. With CO_2_ supplementation, our cultures had glucose-consumption rates of 3.29–3.34 g L^−1^ day^−1^ between cultivation days 1–4 and 0.77–2.21 g L^−1^ d^−1^ without it (Figure 1B, C). This suggests that the presence of CO₂ triggers enhanced glucose uptake and influences central metabolic pathways linked to growth, even in the dark.

The relationship between carbon availability in its native habitats and energy metabolism in *Galdieria* remains an open question. *Galdieria* is found in endolithic and organic-carbon-containing environments in and around hot springs [1,2]. CO_2_ is the major dissolved gas other than N_2_ in sulfate-rich acid hot springs and some dissolved organic matter (DOM) is contributed from surface detritus. DOM is also formed under hydrothermal conditions deep underground, at least in the Yellowstone region of the United States [30], with unknown impacts on *Galdieria* metabolism. Dissolved oxygen will be very low where underground waters reach the surface. Given the low pH and elevated temperatures of hot springs, CO_2_ is expected to outgas as a function of distance from the source while O_2_ will increase with distance. As such, it would not be surprising if *Galdieria* evolved metabolic responses to the transient availabilities of these critical metabolic gases. We sought to investigate this further together with observing culture productivity in different trophic modes with *G. sulphuraria* 5587.1 (Figure 2). This strain was chosen as it shows robust growth behaviors in wastewater-treatment processes [21–23,31].

**Figure 2.**
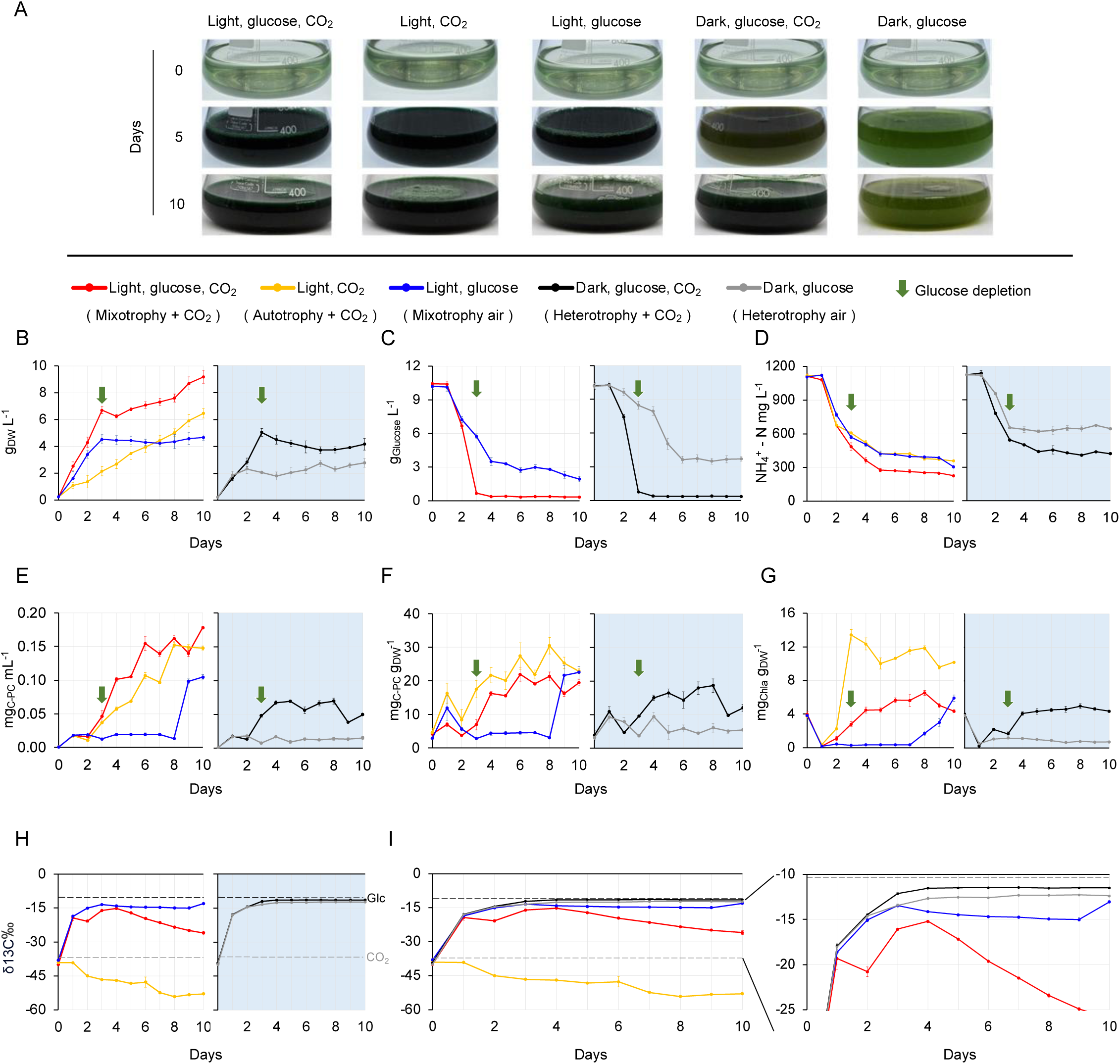
Trophic responses of *G. sulphuraria* 5587.1 to organic carbon and CO_₂_ availability, and their effects on biomass accumulation, pigment content, and isotope composition of biomass. **A**. Representative images of *G. sulphuraria* 5587.1 trophic responses across 10 d of cultivation in 400-mL bioreactors under: (i) mixotrophy with additional 3% CO_₂_ (‘mixotrophy + CO_2_’ - light, glucose, CO_2_), (ii) ‘autotrophy’ with 3% CO_₂_ (light, CO_₂_), (iii) mixotrophy (‘mixotrophy-air’, light, glucose), (iv) heterotrophy with additional CO_₂_ (‘heterotrophy + CO_2_’ - dark, glucose, CO_2_), and (v) heterotrophy without 3% CO_₂_ (‘heterotrophy-air’ - dark, glucose). Cultures were initiated from a single autotrophically grown (12:12 h day:night cycles) inoculum (light, CO_2_). **B–D.** Dry biomass (B), glucose consumption (**C)** Ammonium (NH_₄_^⁺^) consumption (**D)** in cultures grown in the presence of light (left) or in the dark (grey background, right). Arrowheads indicate the day when glucose was depleted in CO_2_-supplemented cultures. **E–G.** Pigment accumulation in biomass over time. **E, F**. C-PC content of the biomass and chlorophyll-a content of the biomass (**G**) in cultures grown in the presence of light (left) or in the dark (grey background, right). **H, I.** Stable-isotope ratios of biomass carbon for – δ^¹³^C (**H)** and overlay of δ^¹³^C values across five conditions for comparison (**I**). Dotted lines mark the δ^¹³^C of the glucose used as a carbon source and the δ^¹³^C of the CO_2_ supplied. At right in (**I**) is a magnified panel view. Error bars indicate the standard deviation of n = 3 replicates.

### Regardless of trophic mode, high CO_2_ promotes *Galdieria* biomass and pigment accumulation

Maintaining photosynthetic capacity while growing on organic carbon should allow an alga to collect photosynthetic electrons, drive the CBB cycle, and fix CO_2_, in addition to the consumption of organic carbon through respiration. It has been proposed that the CO_2_ generated from respiration could be captured by the CBB cycle, when cells are grown with glucose as a carbon source and illumination [20]. Indeed, carbon conversion to biomass yields greater than 0.5 are known for mixotrophic *G. sulphuraria* grown on glucose [26,27]. We wanted to further explore what might cause our observed glucose consumption to be higher with CO_2_ supplementation across *Galdieria*, and so cultured *G. sulphuraria* 5587.1 in five different conditions: complete autotrophy with 3% CO_2_ as a sole carbon source (autotrophy), complete heterotrophy in darkness with 10 g L^−1^ glucose as a sole carbon source (heterotrophy air), heterotrophy with 3% CO_2_ (heterotrophy + CO_2_), illuminated cultures with glucose as a carbon source but no supplemental CO_2_ (mixotrophy air), and illuminated glucose cultures with additional CO_2_ (mixotrophy + CO_2_) (Figure 2A). Cultures were started from the same autotrophically grown preculture as inoculum, from 12 h:12 h day:night cycles with the same light intensity.

Cultures in ‘autotrophy’ proceeded in a linear growth manner with a doubling rate of 0.332 d^−1^ through the 10-d cultivation trial (Figure 2B). This lower growth-rate demonstrates that *Galdieria* are not efficient photosynthetic microbes and may rely more on fermentation/mixotrophy for their lifestyle. Similar to results presented in Figure 1, CO_2_ supplementation had a positive impact on all growth modes, regardless of illumination (Figure 2B). Cultures in ‘heterotrophy + CO_2_’ achieved higher biomass than ‘heterotrophy air’, ‘mixotrophy + CO_2_’ surpassed biomass of ‘mixotrophy air’ cultures (Figure 2B). Glucose was depleted in CO_2_-supplied cultures (heterotrophy + CO_2_ and mixotrophy + CO_2_) by day 3 of cultivation, while non-supplemented cultures (heterotrophy air and mixotrophy air) stalled in glucose consumption at ∼3–4 g L^−1^, concentrations similar to Figure 1B and those reported previously (Figure 2C) [26]. Cultures in mixotrophy + CO_2_ exhibited a peak biomass of ∼6.70 ± 0.27 g L^−1^ at day 3 concomitant with glucose depletion, after which a switch to autotrophy could be observed leading to a final biomass of 9.18 ± 0.49 g L^−1^ on day 10 (Figure 2B). Cultures with mixotrophy air and heterotrophy + CO_2_ both exhibited biomass accumulation peaks of ∼4.66 ± 0.2 g L^−1^ and 4.17 ± 0.44 g L^−1^, respectively, also at day 3, neither increased in biomass after this (Figure 2B). Heterotrophy air ceased biomass accumulation at ∼2.07 ± 0.09 g L^−1^ on day 3, with slow consumption of glucose until stalling and no further ammonium consumption (Figure 2B–D). Growth rates measured for cultures between days 1–3 of cultivation are presented in Table 1.

**Table 1.**
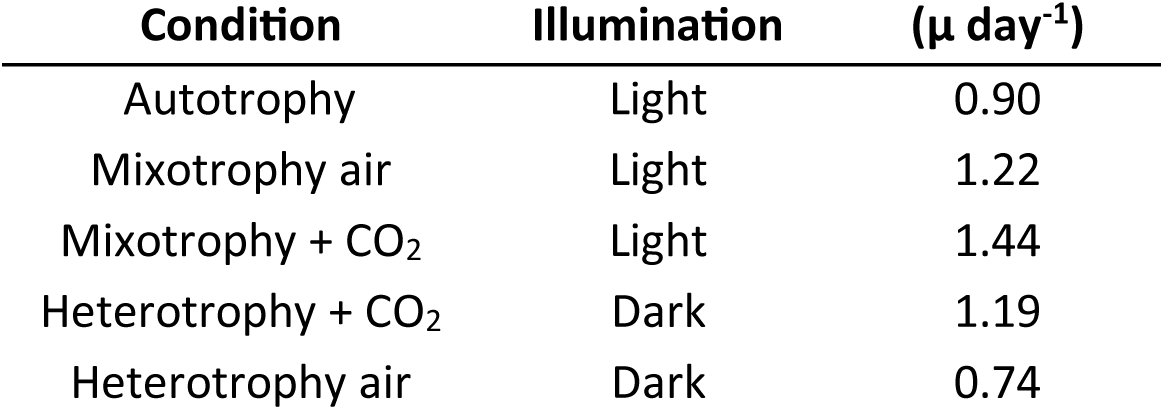
Growth rates (μ, day^⁻¹^) of *G. sulphuraria* 5587.1 cultured under different trophic conditions with or without CO_₂_ supplementation. Growth rates were calculated between days 1– 3 of cultivation.

Ammonium consumption was greater for cultures supplied with CO_2_, except for autotrophic cultures (Figure 2D). Pigment contents were markedly higher in all CO_2_-supplemented cultures (autotrophy, heterotrophy + CO_2_ and mixotrophy + CO_2_) (Figure 2E–G), with the highest C-PC and chlorophyll contents per biomass in autotrophy conditions (Figure 2F,G). Similar trends occurred in heterotrophy + CO_2_ and mixotrophy + CO_2_ regardless of illumination (Figure 2E–G). This was not so in illuminated mixotrophy air, wherein pigmentation was markedly low with a shift occurring at day 8, suggesting a switch to autotrophy (Figure 2E–G). The results indicate dynamic control of intracellular processes in the presence of organic-carbon, and this is triggered by elevated CO_2_ levels.

### Organic carbon, not CO_2_, is consumed in the dark

The culture behaviors described above (Figure 2B–G) suggest CO_2_ plays a role in organic-carbon consumption from the culture medium. Mixotrophy + CO_2_ gave substrate-to-biomass yield ratios of ∼0.7 on day 3 and ∼0.9 on day 10, while both mixotrophy-air and heterotrophy + CO_2_ exhibited a maximum yield of ∼0.5 on day 3 (Figure 2B). The performance differences between heterotrophy-air to heterotrophy + CO_2_ were peculiar, as both cultures would be expected to metabolize glucose equally. The maintenance of pigment content of *G. sulphuraria* 5587.1 in darkness may mean that the CBB cycle is in fact active in heterotrophy, which could explain higher overall biomass in fermentative conditions with elevated CO_2_. However, it is also possible that CO_2_ serves as a metabolic trigger, stimulating heterotrophic carbon metabolism by an unknown cellular mechanism. To disentangle the nuances of this relationship, stable-isotope analysis was used to determine the role of each carbon source in *Galdieria* biomass formation.

Carbon-isotope ratios (δ^13^C) accurately reflect the carbon source used in photosynthetic carbon fixation or mixotrophic biomass production [32]. These ratios are commonly used in ecological studies to identify the primary carbon sources at the base of food webs [33], and δ¹³C values offer insights into their biological and geological origins and can have slight variations over short timescales. For example, C₃ plants, which rely on the CBB cycle for carbon fixation, typically exhibit δ¹³C values between −20‰ and −34‰, with an average of around −27‰. In contrast, C₄ plants, which follow a different photosynthetic pathway, tend to have δ¹³C values ranging from - 9‰ to −16‰, averaging near −13‰ [34]. Fossil fuels, such as petroleum, display broader δ¹³C ranges, from approximately −44‰ to −19‰, reflecting variations in their biological sources and geological history [35]. We applied this technique to *Galdiera,* for the first time reported in literature, to tease apart its mixotrophic and heterotrophic responses to glucose and CO_2_. Daily biomass samples from the bioreactor experiment in Figure 2 were analyzed for δ^13^C ratios at each time point (Figure 2H,I, Supplemental Figure 2D). If cells consume organic carbon from the medium, biomass would be enriched in ^13^C and trend towards values of the glucose standard (δ^13^C = –10.53 ± 0.12‰, Figure 2H). However, if the supplied CO_2_ was taken as a carbon source, biomass δ^13^C values would have ^13^C reflecting more negative values like that of CO_2_ (δ^13^C = –36.29 ± 0.27‰, Figure 2H). The carbon source consumed in biomass formation was clearly represented in the δ^13^C values of *Galdieria* biomass (Figure 2H, I).

Autotrophic cultures exhibited depleted (negative) δ^13^C values that started from –35‰ and trended to –50‰, with fossil-derived CO_2_ the sole carbon source. Mixotrophy + air cultures were enriched in ^13^C (trending towards positive) with δ^13^C values starting from –35‰ and trending to –15‰, indicating glucose consumed as the carbon source (Figure 2H, I). In the dark, δ^13^C values for heterotrophy + CO_2_ were higher than heterotrophy air (Figure 2H, I). This indicates that glucose is more effectively consumed by heterotrophy + CO_2_ cultures than in the heterotrophy air condition, as the opposite would be true if the enriched CO_2_ was being consumed (Figure 2I). This strongly suggests that CO_2_ is acting as a modulator of metabolism without direct involvement of the CBB cycle.

In mixotrophy air, glucose consumption is inefficient (Figure 2C), with δ^13^C values suggesting either a contribution of respired CO_2_ or loss of organic carbon through respiration after day 3 (Figure 2H,I). Until this point, δ^13^C values are similar in mixotrophy air to that of heterotrophy air (Fig 2M), after which growth was arrested (Figure 2B). It could be that respiratory CO_2_ is consumed in the CBB cycle, however, the slow metabolism of glucose from the medium indicates that respiration is active, but not sufficient, to stimulate faster growth and glucose consumption (Figure 2C). This again suggests a high-CO_2_ environment is required to cue the cell into active growth, but the mechanism of this is unclear. In mixotrophy + CO_2_, a slightly lower δ^13^C value is observed compared to other glucose-containing conditions until day 4, after which a linear reduction occurs following what likely is a shift to autotrophy after glucose consumption (Figure 2H,I). This lower δ^13^C value supports the notion that in illuminated mixotrophic conditions, supplemental CO_2_ is consumed by the cells in addition to the organic glucose carbon source, leading to the observed higher biomass yields (Figure 2B).

Our results suggest that high CO_2_ stimulates active metabolism that leads to efficient organic-carbon uptake in a way that is not yet understood. Specifically, high CO_2_ levels in fermentative conditions lead to complete metabolism of glucose from the medium, whereas a low-CO_2_ environment results in stalled fermentation. Only under illuminated conditions is CO_2_ consumed in a small amount, concomitant with organic-carbon consumption, or autotrophy. We have previously observed that smaller shake flasks growing mixotrophically can effectively consume organic carbon (not shown) and a previous report with significantly higher glucose supplementation rates but no supplemental CO_2_ also showed efficient heterotrophy [13]. We speculate that if cells can consume enough glucose in the early days of cultivation, they could generate their own CO_2_ atmosphere through respiration, which then stimulates complete consumption of glucose through the CO_2_-induced signal. This response of the cells to high CO_2_ may have confounded previous analysis of their growth behaviors and may not have been noticed in lab-scale analysis. The use of headspace recycling [26,27] likely supports maintenance of this favored condition and the high biomass yields observed in oxygen-balanced mixotrophy. Our findings indicate the requirement of high CO_2_ is essential for both fermentative and mixotrophic cultivation of *Galdieria*, at least under the tested conditions.

### *Galdieria* consumes many carbon sources, but weak acids killed the extremophile

*Galdieria* is reported to consume over 50 known carbon sources [7,36–38], however, our analysis of peer-reviewed studies found only 34 (Table 2). *Galdieria* reportedly does not grow on many organic acids, with only one report showing this for very low concentrations of acetic acid [18]. A goal of working with *Galdieria* bioproduction is the conversion of waste carbon into biomass. Many mixed organic-carbon wastes will have organic acids within them, such as those from food manufacture or discarded food wastes. We subjected *G. sulphuraria* 5587.1 to different concentrations of seven weak organic acids to determine if the cells would use them as carbon sources or tolerate them (Supplemental Figure 1).

**Table 2.** Carbon sources used for growth in *Galdieria* in previous studies.

| Carbon Source | Molarity tested mM or g L<sup>-1</sup> | Reference |
| --- | --- | --- |
| Sucrose | 12.5–277.5 | [6][9] |
| D-Glucose | 25 | [6][8] |
| L-Glucose | 25 | [6] |
| D-Mannose | 25 | [6] [8] |
| D-Galactose | 25–69.4 | [6][7] |
| D-Fructose | 25 | [6] [8] |
| L-Sorbose | 25 | [6] [8] |
| D-Fucose | 25 | [6] [8] |
| L-Fucose | 25 | [6] [8] |
| L-Rhamnose | 25 | [6] [8] |
| 2-Deoxy-D-galactose | 25 | [6] |
| D-Arabinose | 25 | [6] |
| L-Arabinose | 25 | [6] |
| D-Lyxose | 25 | [6] |
| D-Ribose | 25 | [6] [8] |
| D-Xylose | 25 | [6] [8] |
| L-Xylose | 25 | [6] |
| D-Sorbitol | 25 | [6] [8] |
| D-Mannitol | 25 | [6] [8] |
| Xylitol | 25 | [6] [8] |
| L-Arabitol | 25 | [6] [8] |
| D-Arabitol | 25 | [6] [8] |
| Adonitol | 25 | [6] [8] |
| Glycerol | 25 | [6] |
| Erythritol | 25 | [6] |
| Dulcitol | 25 | [6] [8] |
| L-Fucitol | 25 | [6] |
| Meso-erythriol | 25 | [6] |
| Myo-inositol | 25 | [6] |
| 6-Deoxy-glucose | 25 | [8] |
| Maltodextrin (Paselli-SA2) | 10–50 g L<sup>-1</sup> | [9] |
| Granular starch<br>(potato and corn) | 10 g L<sup>-1</sup> | [9] |
| Lactose | 34.7 | [7] |
| Acetic acid | 8.33 | [39] |
| Chocolate premix | 30 g L<sup>-1</sup> | This study |

Formic, acetic, oxalic, lactic, succinic, and tartaric acids were all lethal or inhibitory to *Galdieria* at 10 mM (0.46–1.5 g L^-1^). Serial attempts to adapt strains to higher concentrations were unsuccessful (not shown). Citric acid was tolerated by *G. sulphuraria* 5587.1 regardless of concentration (Supplemental Figure 1). These data suggest that small-molecule weak acids are all toxic to the acidophile, thereby limiting the types of feedstocks usable in its cultivation. This is likely due to the ability of such acids to cross biological membranes, being fully protonated in the low-pH environment outside the cell and deprotonating inside the neutral cytosol.

### Confectionary-production waste as substrate for *Galdieria* biomass production

Although each confectionary recipe is unique, cacao, butter, and sugar are mixed prior to a high heat pasteurization treatment to make chocolate bars. This substance, hereafter referred to as ‘chocolate premix’, is high in caloric value. During typical factory-operation cycles, or when production ceases, process lines start and stop prior to pasteurization, causing losses of chocolate premix, around 20 metric tons annually for a mid-sized factory (Mars facility, King Abdullah Economic City (KAEC), Saudi Arabia, personal communication). We acquired samples of waste chocolate premix from a local confectionary factory and investigated whether it could be revalorized into *G. sulphuraria* biomass (Figure 3A, B). This substrate can be consumed as a carbon source by *G. sulphuraria* in bioreactors (Figure 3A, B). The nuances of that consumption are explored in the following sub-sections.

**Figure 3.**
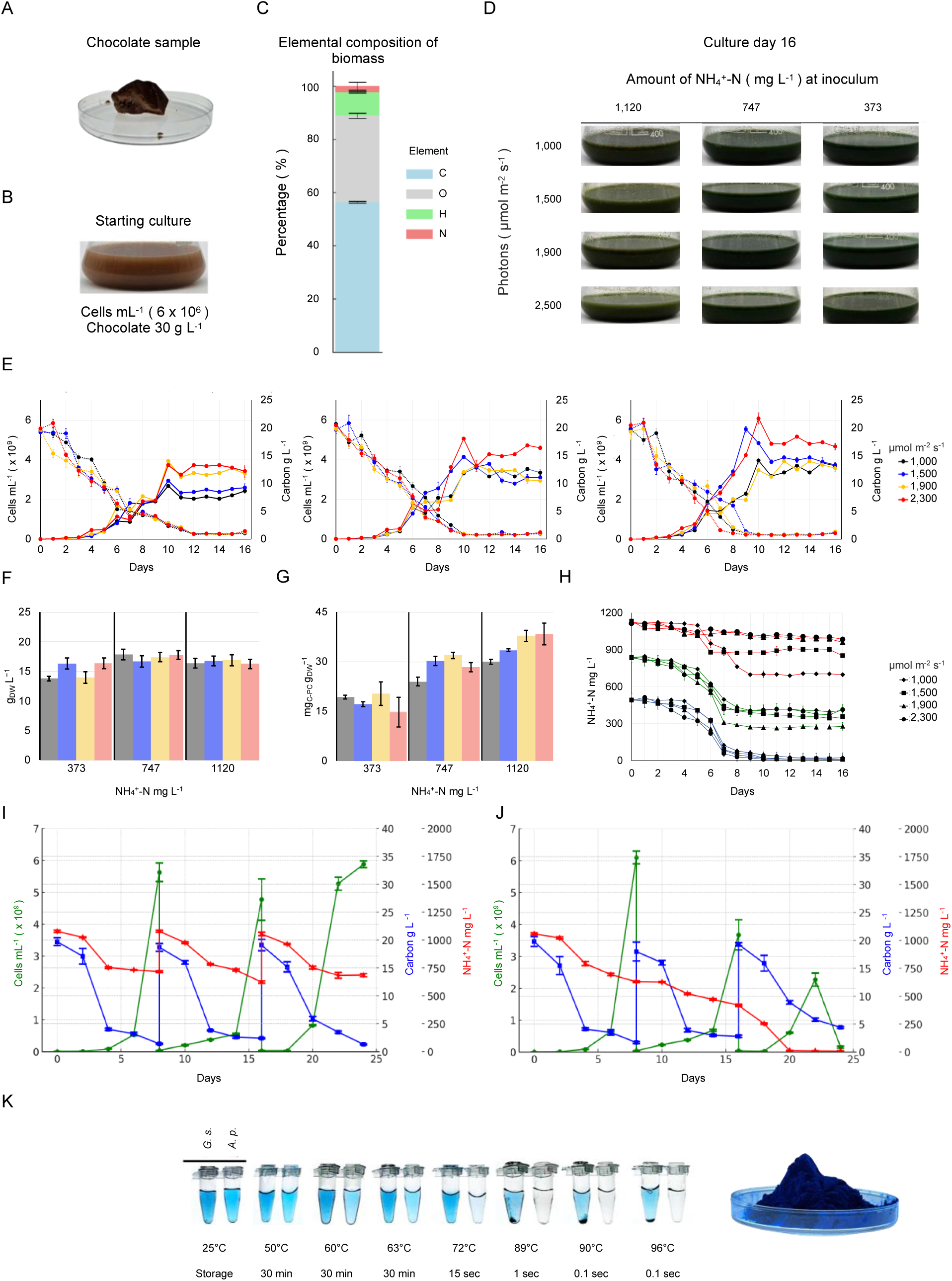
Growth of *G. sulphuraria* 5587.1 on chocolate manufacture waste and thermostability of C-phycocyanin. **A.** Photograph of a solid chocolate waste sample. **B.** *G. sulphuraria* culture initiated with 30 g L^⁻¹^ chocolate waste as the sole organic carbon source. **C.** Elemental composition of chocolate waste. **D.** Cultures after 16 d of growth under varying NH_₄_^⁺^-N concentrations and light intensities. **E.** Growth dynamics of cultures from (D). Left: Culture with 373; middle: 747 and right: 1,120 mg NH_₄_^⁺^-N l^⁻¹^. **F.** Average biomass accumulation over the final 3 d of cultivation. Bars represent mean biomass values from the last 3 days, with colors corresponding to different light intensities. **G.** Average C-PC production over 3 d of cultivation. Bars represent mean C-PC content per biomass dry weight from the last three days, with colors indicating different light intensities. **H.** Ammonium (NH_₄_^⁺^-N) consumption from cultivation experiments in (D). The colors indicate the initial NH_₄_^⁺^-N concentration (red: 1,120, green: 747, blue: 373 mg L^⁻¹^). **I, J.** Growth during cyclic reuse of chocolate culture medium for continuous biomass production. **I**: Filtered medium replenished with both NH_₄_^⁺^-N and chocolate waste every 8 d for three cycles. **J**: Filtered medium replenished only with chocolate waste. **K.** Left: Thermostability comparison of *G. sulphuraria* 5587.1 and *Limnospira* (*Spirulina*) C-PC extracts. Right: Freeze-dried C-PC extract from *G. sulphuraria* 5587.1.

### High ammonia is required for efficient carbon consumption and pigment accumulation

Sampled chocolate mass contains very low nitrogen, but over 57% carbon (Figure 3C). To use this chocolate as a substrate for growth, we generated culture medium with chemically supplied ammonium as a nitrogen source and enough chocolate to supply 20 g L^−1^ organic carbon in addition to supplemental 3% CO_2_ in 400-mL photobioreactors as above (Figure 3B, D). Variations in nitrogen and light intensity across a range of concentrations were investigated. For all cultures, light intensity did not influence the rate of organic carbon consumption (Figure 3E). Cultures with the highest nitrogen content exhibited total carbon consumption by ∼day 8, with increasing time to complete consumption with decreasing NH_4_ concentrations (Figure 3E). Glucose-uptake rates were directly related to higher NH_4_ contents, at 1,120 mg L^−1^ (same as MA2 medium) glucose was consumed at an average 3.06 g L^−1^ d^−1^, while 747 mg L^−1^ the rate was 2.31 g L^−1^ d^−1^, and 373 mg L^−1^ down to 2.21 g L^−1^ d^−1^ between days 1–4. Light intensities above 1,900 μmol photons m⁻² s^⁻¹^ gave higher final cell counts than 1,500 μmol photons m⁻² s^⁻¹^ with divergence around the time points when organic carbon drops below 5 g L^−1^ (Figure 3E). These results mirror previous findings that lower C:N ratios resulted in more efficient batch production of *Galdieria* in mixotrophy or heterotrophy [14].

Total biomass could be quantified only in the last days of cultivation due to the presence of chocolate in the culture medium in earlier time points, however, the final biomass amounts in each cultivation were not different across nitrogen contents or light intensities (Figure 3F). However, C-PC content trended higher under the highest nitrogen contents and light intensities (Figure 3G). Final C-PC contents across conditions ranged from 14.76 ± 4.46 to 38.4 ± 3.31 mg g^−1^ DBM (1.47–3.84%) (Figure 3G). Although these yields are lower compared to previously reported values of 13.88% [15], they are consistent with other studies [13,14,40]. Nitrogen consumption was consistent within starting concentration groups, with higher light intensities seemingly showing lower rates of nitrogen consumption in the highest nitrogen-containing group (Figure 3H). The results indicate that in addition to requiring high CO_2_, *Galdieria* also require high nitrogen contents in its surroundings to stimulate efficient organic carbon consumption. This is in line with previous observations [7,21]. Likely, C-PC yields and carbon-consumption rates can be increased further by using lower C:N ratios in mixotrophic set-ups. However, the high ammonium concentration of the medium at the end of cultivation presents a challenge, as it requires removal from the liquid before discharge.

### Cyclic reuse of culture medium enables safe conversion of carbon waste to biomass

The requirement for high ammonium concentrations to drive mixotrophic biomass accumulation in *Galdieria* means that waste-valorization concepts necessitate review of bioprocess planning to prevent high ammonium-containing effluent discharge at the end of cultivation. Ammonium was not fully consumed during chocolate-waste mixotrophy (Figure 3H). Therefore, we attempted to reuse the culture medium and determine if any growth inhibition could be observed in repetitive uses (Figure 3I, J). After ∼8 d of cultivation, ∼70% of the culture was harvested, biomass separated, the medium subjected to ultrafiltration, and either refreshed with ammonium and chocolate waste, or only chocolate waste. Three repetitive cycles showed no decline in culture growth for medium refreshed with ammonium (Figure 3I), while omitting nitrogen allowed its complete consumption within 20 d, but graded reduction in *Galdieria* growth was observed (Figure 3J). These findings indicate it should be possible to run cyclic processes with *Galdieria*, maintaining strict control of water reuse and ammonium doping to effectively turn organic-carbon wastes into algal biomass.

### *Galdieria* phycocyanin is pasteurizable

The blue photopigment C-PC is a water-soluble protein complex forming part of the alga’s light-harvesting machinery [41]. To extract C-PC, cell lysis and separation of total soluble protein is usually performed, generating a blue liquid with proteins containing C-PC [42]. This can be dried and used as a food coloring or pigment, among other applications in medical imaging [43–45]. Commercially available C-PC is sourced from the cyanobacteria collectively called ‘Spirulina’ [46] within the genus *Limnospira* (formerly *Arthrospira*) [47]. C-PC from *Limnospira* is not thermotolerant, which challenges its pasteurization and broader applicability [48]. Other evidence of C-PC in Galdieria shows that C-PC from *G. phlegrea* also exhibits thermostability around 70 °C and its production varies depending on its trophic condition [12]. Here, we tested C-PC extracts from *G. sulphuraria* 5587.1 against that of commercial *Limnospira* across a range of pasteurization treatments (Figure 3K). C-PC from *G. sulphuraria* 5587.1 could withstand 72°C for 15 s without reduction in color intensity or precipitation, whereas the intensity of *Limnospira* C-PC reduced already at 63°C for 30 min (Figure 3K). The results suggest that in addition to the potential to produce a protein-rich biomass from waste carbon inputs that contain a complete amino-acid profile, *G. sulphuraria* could also generate thermostable C-PC as a co-product of its biomass fractionation.

Our findings suggest that *G sulphuraria* 5587.1 can efficiently transform industrial chocolate waste into high-value biomass and phycocyanin as a by-product. In batch cultures with 30 g L^⁻¹^ chocolate waste, *G. sulphuraria* 5587.1 produced up to 17.85 g L^⁻¹^ biomass and 686 mg L^⁻¹^ C-phycocyanin within 10 d, corresponding to ∼0.595 g biomass and 0.0229 g C-PC per g chocolate. Under batch-fed conditions with three 8-d cycles, cumulative yields were 53.55 g L^⁻¹^ biomass and 2.06 g L^⁻¹^ C-PC, improving daily productivity to 2.23 g L^⁻¹^ d^⁻¹^ and 86 mg L^⁻¹^ d^⁻¹^, respectively. A medium-sized chocolate factory in Saudi Arabia generates approximately 20 metric tons of this waste annually (personal communication), which, if fully used under these conditions, can yield ∼11.9 metric tons of algal biomass containing ∼457 kg C-PC. The extremophilic growth conditions of *Galdieria*—low pH (1–2.5) and high temperature—can help minimize microbial contamination, with less need for sterility. Further, external industrial CO₂ can be used as a carbon source. These findings show the potential for large-scale algal cultivation to valorize industrial wastes into value-added algal bioproducts and contribute to circular-bioeconomy and sustainability goals. Together, these advantages of low-cost inputs in the form of chocolate waste or other similar feedstocks are appealing as favorable substrates for cultivation of this extremophilic genus. This work provides a framework for repurposing confectionery by-products into valuable algal-derived bioproducts, supporting waste valorization and circular-bioeconomy strategies in the region.

### Conclusions

Since its discovery, members of *Galdieria* have intrigued biotechnologists and phycologists alike. Its extreme habitat is at the limits of eukaryotic temperature and physiological pH tolerances at the interface with geochemical processes in a way that most mesophiles are not. Its genetics are supported by a large component of horizontally transferred genes [49], which may have assisted in its adaptation to this unique lifestyle and broad carbon-substrate capacity. *Galdieria* have been proposed for use in fermentative bioprocesses to convert low-value, or waste, carbon sources to higher-value algal biomass for either generation of C-PC or the protein-rich biomass itself. Besides confectionery applications, *G. sulphuraria* 5587.1 is promising as a single-cell protein, nutraceutical, and natural food colorant due to thermostability of its phycocyanin, antioxidant potential, and quality amino acid composition of its biomass. Its ability to grow on diverse substrates such as glycerol, xylitol, glucose, and industrial food wastes supports their valorization in circular economies. This strain also possesses the ability to accumulate C-PC under dark, CO_₂_-enriched, glucose-fed conditions. These findings support a low-cost, scalable, heterotrophic bioprocess for protein and pigment production irrespective of photosynthesis, which support continuous, indoor, light-independent production, unlike conventional light-dependent systems, such as *Limnospira*.

Our observations have revealed a previously overlooked feature of its metabolism, that elevated CO_2_ seems to trigger the cell into an active metabolic state to consume external organic carbon. It is likely that in small-scale shake flasks or even bioreactor fermentation, a CO_2_ atmosphere can form from respiration in the early stages, leading to continued active metabolic states. Indeed, fermentations have been successfully conducted with the organism since its discovery [13]. We were surprised by the differential response of *Galdieria* based on CO_2_ presence across all tested strains and in darkness as observed here. The biological mechanism which CO_2_ stimulates remains the subject of further analysis and may be revealed in future studies via transcriptomics and proteomics. Stable-isotope analysis showed that, indeed, CO_2_ is stimulating active metabolism but is not being consumed in darkness, while CO_2_ can be consumed in illuminated mixotrophy but at lower amounts than glucose. Our findings will influence future efforts with this organism, which could either be maintained in closed atmospheres, or a high-CO_2_ mixture in gas sparging to promote this response and achieve higher biomass. Despite being able to tolerate extreme conditions, weak acids have a detrimental effect on *Galdieria*, presumably via uptake in the protonated form followed by deprotonation in the cytoplasm. This means that waste carbon substrates must be appropriately chosen as feedstocks, as those high in organic acids will not be suitable. The positive side to this, however, is that readily available substances like acetic acid, can be used as a biocontainment agent in its cultivation facilities.

## Acknowledgments

We are grateful to Mustafa Bin Kamaludin, Mohamed Abo El-Ola, and Frank Mars for facilitating access to the KAEC Mars facility in December 2023 and providing chocolate-waste samples. We thank Naydu Zambrano from KAUST for the support with elemental analysis of the chocolate samples. *G. partita* NBRC102759 was obtained from Shin-Ya Miyagishima. KJL and MDF acknowledge baseline research funding from KAUST. MLPM is funded by the KAUST opportunity fund grant #5576 awarded to KJL. PJL acknowledges funding from ASU Lightworks supporting Cyanidiophyceae research.

## Materials and Methods

### Algae cultures and growth tests

Strains of *G. sulphuraria* were obtained from the following sources: CCMEE SOOS (from A. Weber, Heinrich Henne University), 5587.1 and 5572 were obtained from R. Castenholz (University of Oregon, [1]). Strains 074G, 21.92 and *G. javensis* 074W, reclassified from *G. sulphuraria* 074W, were from the SAG culture collection (Götinggen, Germany). *G. sulphuraria* 3377 was isolated in the Lammers’ lab via colony formation on glucose plates using an inoculum of *C. merolae* 10D obtained directly from NIES culture collection (Japan). *G. partita* NBRC102759 was obtained from NIES collection from Shin-Ya Miyagishima.

Cells were routinely cultured in MA2 liquid medium [50] at pH 2.5, adjusted with H_₂_SO_₄_, in 50-mL flasks shaken at 100 rpm under continuous white light (90 μmol photons m^⁻²^ s^⁻¹^) at 42°C in a Percival incubator (AL-30, Percival Scientific), supplemented with 3% CO_₂_ in air mixtures. For heterotrophic and mixotrophic cultures, 10 g L^⁻1^ glucose was added by filter sterilization after autoclaving.

All precultures for experiments were performed by inoculating 1 x 10^6^ cells mL^⁻1^ in a 50-mL working volume of MA2 within 125-mL Erlenmeyer flasks with 110 rpm shaking for 96 h. Cells were then resuspended to the desired densities in 100-mL working volumes in 400-mL Erlenmeyer flasks before growth analysis, which was performed in a 400-mL working volume in 1-L Erlenmeyer flasks and grown in Algem photobioreactors (Algenuity©, United Kingdom). Growth experiments were carried out under different conditions (Table 3) in photobioreactors, except in experiments where CO_₂_ was excluded. Continuous illumination was provided at 1,000 μmol photons m^⁻²^ s^⁻¹^ for autotrophic and mixotrophic cultures. Light shades were used to block light entry completely for heterotrophic cultures. Spectra of the bioreactor light-emitting diodes can be found in the Supplement of [51].

**Table 3.** The eight combinations of light, glucose and enriched CO_₂_ were tested for the growth of *G. sulphuraria* 5587.1 as presented in Figure 2. The presence or absence of each variable is indicated for each experiment.

| Condition | Light<br>(1,000 photons m<sup>-2</sup> s<sup>-1</sup>) | Glucose (10 g L<sup>-1</sup>) | CO<sub>2</sub> (3%) |
| --- | --- | --- | --- |
| 1 | + | + | + |
| 2 | + | + | - |
| 3 | + | - | + |
| 4 | + | - | - |
| 5 | - | + | + |
| 6 | - | + | - |
| 7 | - | - | + |
| 8 | - | - | - |

Chocolate mass, butter and cacao solid, collectively ‘chocolate waste’, discarded during confection production was used as a substrate for *G. sulphuraria* 5587.1 growth. Chocolate waste was collected at the Mars confectionary factory, King Abdullah Economic City, Saudi Arabia in December 2023 and stored at 4°C until use. To prepare the culture medium, 30 g L⁻¹ of chocolate sample was weighed and dissolved in sterile distilled water (ddH_₂_O) by continuous stirring at 50°C and the addition of ammonia as indicated.

To assess the feasibility of fed-batch cultures with chocolate waste as the primary substrate, *G. sulphuraria* 5587.1 was acclimated to the medium before starting repetitive cultivation operations. Acclimation was achieved by culturing the strain on the chocolate-waste media for 2 w, after which the culture was used to inoculate freshly prepared medium using chocolate as substrate.

For repetitive cultivations, after the depletion of the carbon source, cultures were centrifuged at 2,000 x *g* for 10 min, and the supernatant was filtered through a 0.45-μm filter. Subsequently, 30 g L^⁻¹^ of fresh chocolate sample was added, and the ammonia content was regularly monitored and replenished as indicated throughout the experiment. Cells (1 × 10^⁷^) were reintroduced into the culture. The experimental conditions applied to the chocolate waste cultures are described in Table 4. Data for all figures can be found in Supplemental Data File 1.

**Table 4.**
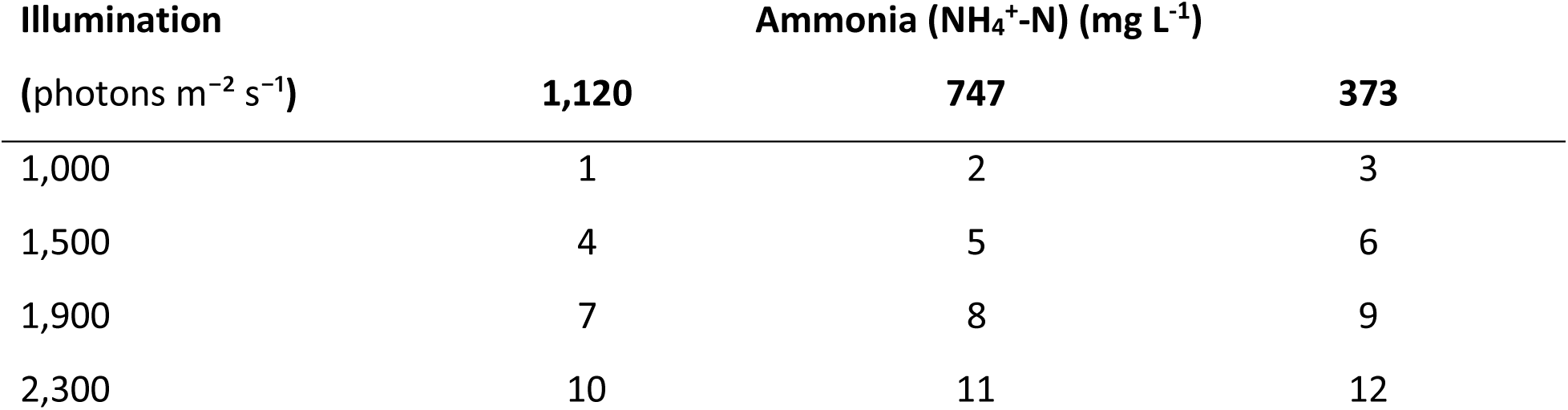
Ammonia concentration and light-intensity combinations used in the experiment with *G. sulphuraria* 5587.1 cultured in chocolate waste presented in Figure 3. Specific combinations of ammonia and light intensities were investigated. Numbers represent flasks listed in Figure 3.

### Phycocyanin extraction

Phycocyanin (C-PC) from *G. sulphuraria* 5587.1 was extracted by a bead-beating FastPrep-24 5G^™^ (MP Biomedicals). For each sample, 200 mg of 1-mm diameter glass beads were added to the concentrated wet biomass with 1 mL of PBS buffer (pH 7.4), precooled to 4°C, and ran for 60 s at 6 m s^−1^ for 3 cycles with 60 s of intermediate cooling. Each sample was then centrifuged at 10,000 x *g* for 10 min. To measure the C-PC content, each sample was measured against a buffer blank at 615 nm (A_615_) and 652 nm (A_652_), respectively, and the absorbance of the cellular debris at 720 nm (A_720_) in a UV–Vis Spectrophotometer (Thermo Scientific® Genesys™ 50, United Kingdom). The result was used to calculate the concentration of C-PC according to Equation (1) [52]:

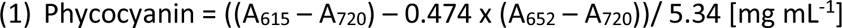

### Chlorophyll a content

Pigments from *G. sulphuraria* 5587.1 were extracted following the same bead-beating protocol used above but with acetone:methanol:water (90:5:5), and absorbance was measured at 630 nm (A_630_), 647 nm (A_647_), and 630 nm (A_630_). The result was used to calculate chlorophyll a concentration according to Equation (2) [53].

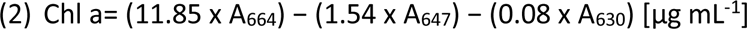

### Chlorophyll-fluorescence measurements

Chlorophyll-fluorescence parameters were assessed using a pulse amplitude modulation (PAM) fluorometer (Mini-PAM-II, Heinz Walz GmbH, Germany). Prior to data acquisition, the signal amplitude was optimized, and three single-turnover measurements (n = 3) were subsequently recorded for each algae sample [54].

### Culture growth monitoring

Dry cell weight was measured by centrifuging 3 mL of culture medium at 2,000 x *g* for 10 min using 13-mL glass tubes. The supernatant was discarded and the remaining biomass was dried at 100°C for 24 h. Cell density was measured using a Neubauer hemocytometer and a 10-μL aliquot of well-mixed culture was pipetted onto the hemocytometer, and cells were counted under a light microscope at 40x magnification. The average cell count from four large squares of the grid was used to calculate the cell concentration according to the formula:

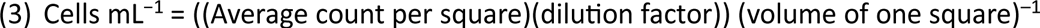

### Nutrient contents

Glucose concentration was measured using the phenol–sulfuric acid method, with a standard glucose curve (from 0–200 μg mL^−1^) [55]. A fixed volume of culture (1 mL) was collected and appropriately diluted as needed to fit within the range of the standard curve. For each sample, 1 mL of 5% (v/v) phenol solution was added, followed by the addition of 5 mL of concentrated H_2_SO_4_. The reaction was mixed gently and allowed to stand at room temperature for 10 min. After the initial reaction, samples were left to cool in an ice bath for 30 min. The absorbance was measured at 490 nm using a UV–Vis Spectrophotometer (Thermo Scientific® Genesys™ 50, United Kingdom), and the carbon concentration of each sample was determined by comparing the absorbance values to the standard curve. Results were expressed as g L^−1^.

Nitrogen concentrations in the culture medium at different time points were determined as follows: 1 mL of each algae culture was sampled daily, centrifuged at 2,000 x *g* for 10 min at room temperature, and the supernatant collected. Ammonium (NH_4_^+^-N) concentration was determined spectrophotometrically with a DR 1900 spectrophotometer (Hach, Germany) after the samples were prepared with AmVer High Range analysis kits (Hach, USA), following the manufacturer’s protocols. Each ammonium concentration was analyzed in technical triplicates.

### Elemental analysis

The elemental analysis (C, H, N, O and S) of 10 mg of lyophilized waste chocolate mass was performed using a Flash 2000 CHNS/O analyzer (Thermo Fisher Scientific, Germany). 2,5-(Bis(5-tert-butyl-2-benzo-oxazol-2-yl) thiophene (C: 72.53 wt%, H: 6.09 wt%, N: 6.51 wt%, O: 4.73 wt%) was used as a standard to build the calibration curve, and all samples were measured in triplicate to ensure repeatability.

### C-PC pasteurization

Phycocyanin was extracted from *G. sulphuraria* 5587.1 as above, followed by lyophilization and resuspension in distilled water. Commercially sourced C-PC from *Limnospira* (formerly *Arthrospira*) *platensis* was also resuspended in distilled water. For pasteurization, 50-μL aliquots of 0.2 mg mL^−1^ of each phycocyanin sample were prepared and subjected to each different temperature treatment. The temperature was maintained using a T100 Thermal Cycler (Bio-Rad) to ensure precise and consistent heating for each experiment.

### Carbon and nitrogen stable-isotope analysis (δ^13^C and δ^15^N)

Samples from each experimental treatment (n = 3 replicates at treatment level) were measured in triplicate using a ThermoFisher Scientific™ (Germany) DELTA™ Q Isotope Ratio Mass Spectrometer coupled with an EA – FLASH, IsoLink CN Elemental Analyzer. Isotopic values are reported as delta (δ) values where δ = 1,000 × [(R_sample_/R_standard_) − 1] and R_sample_ or R_standard_ are expressed as the ratio of the heavy to light isotope in per mil (‰) relative to the international standards, Vienna Pee Dee Belemnite (V-PDB) for δ^13^C and atmospheric N_2_ for δ^15^N. To accommodate the analysis of gaseous CO_₂_, modifications were made to the standard protocol. As a solid form of the CO_₂_ sample was unavailable, direct injection was performed using a 5 mL gas-tight syringe (Restek Corporation, USA) into the continuous-flow interface. Scale normalization was performed using two urea working standards (δ^¹³^C –35.46‰ and –42.18‰), USGS44 (δ^¹³^C –42.21‰), and a certified CO_₂_ reference gas (Airgas, Inc., USA) with a known δ¹³C value of –2.7 ± 0.5‰ traceable to VPDB scale. Due to the depleted δ^13^C values of *G. sulphuraria* 5587.1 grown under 3% CO_2_ in the absence of glucose (δ^13^C –47.8 ± 5.3 ‰), we used USGS44 (certified δ^13^C = –42.21‰) as a quality-control standard (SD of replicate measurements –42.21‰ ± 0.05). Repeated measurements of internal reference material calibrated against IAEA600, USGS64, and USGS40, Cat No. B2174 produced an average within run precision (SD) of 0.1‰ for δ^13^C and 0.2 ‰ for δ^15^N. In this study, the used carbon sources were characterized by their δ^¹³^C signatures: the supplemented CO_₂_ exhibited a δ^¹³^C value of −36.29 ± 0.27‰, while the supplemented dextrose had a δ^¹³^C value of −10.53 ± 0.12‰.

