## Supplemental Figures 1&2 for "Elevated carbon dioxide stimulates highly efficient organic-carbon consumption and confectionary-waste valorization under mixotrophy in the unicellular alga *Galdieria*"

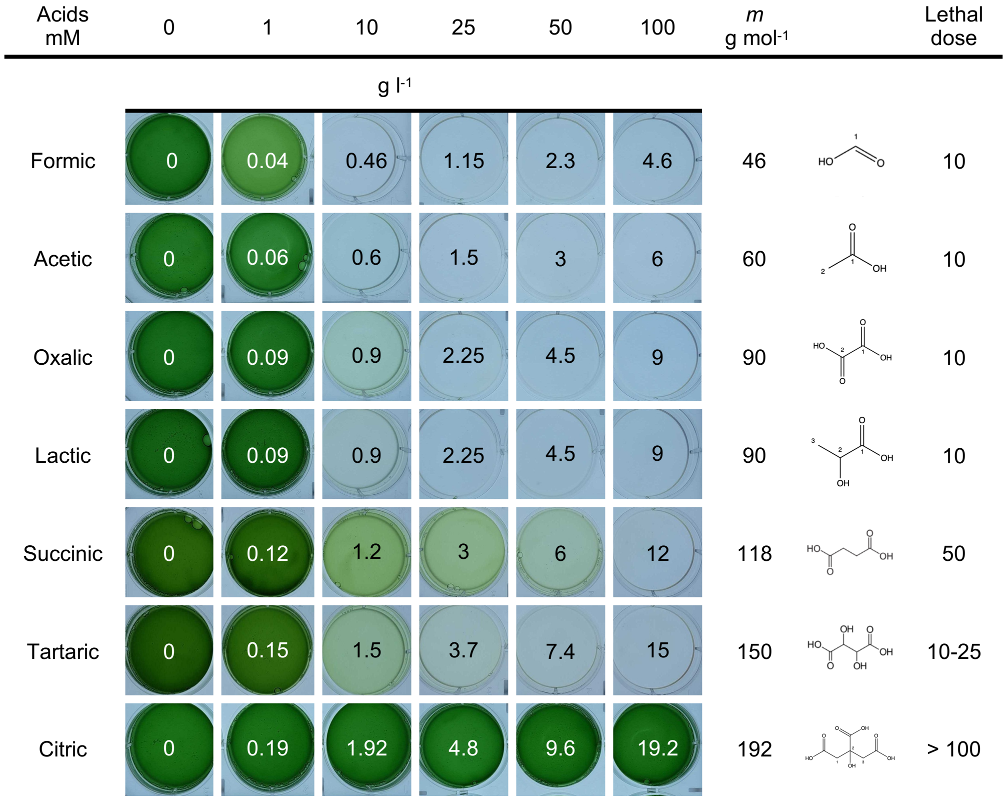


**Supplemental Figure 1. *G. sulphuraria* tolerance to small molecule weak organic acids.** Growth of *G. sulphuraria* in MA2G medium supplemented with increasing concentrations of various organic acids in 5-mL wells. The culture pictures indicate the concentrations (g $L^{-1}$) and the molecular weight (g $mol^{-1}$) of each organic acid is listed alongside its molecular structure. The lethal dose represents the concentration at which no growth was observed.

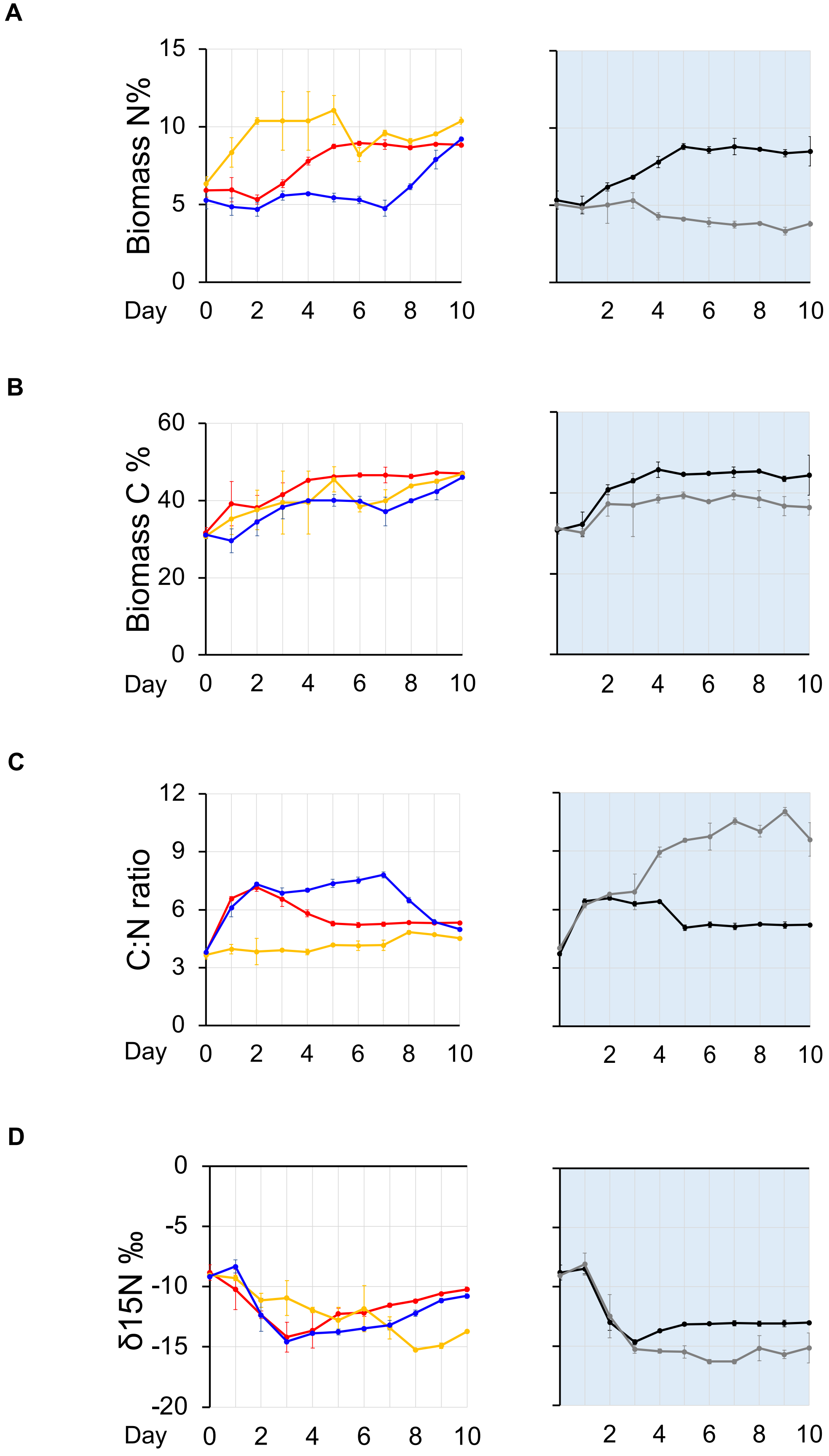


**Supplementary Figure 2. Trophic responses *G. sulphuraria* 5587.1 to organic carbon and $CO_2$ availability, and their effects biomass and isotope composition. A–C.** Elemental composition of biomass. **A:** Biomass nitrogen content (%N). **B**: Biomass carbon content (%C). **C:** Carbon-to-nitrogen (C:N) ratio. **D**. Stable-isotope ratios of biomass nitrogen in $\delta^{15}N$ values. Error bars indicate standard deviation of n = 3 replicates.
